# Discovery of the missing cytochrome P450 monooxygenase cyclases that conclude glyceollin biosynthesis in soybean

**DOI:** 10.1101/2024.07.04.602010

**Authors:** Praveen Khatri, Kuflom Kuflu, Tim McDowell, Jie Lin, Nikola Kovinich, Sangeeta Dhaubhadel

## Abstract

Glyceollins are isoflavonoid-derived metabolites produced by soybean that hold great promise in improving human and animal health due to their antimicrobial, and other medicinal properties. They play important roles in agriculture by defending soybean against one of its most destructive pathogens, *Phytophthora sojae*. Longstanding research efforts have focused on improving accessibility to glyceollins, yet chemical synthesis remains uneconomical. The fact that some of the key genes involved in the final step of glyceollin biosynthesis have not been identified, engineering the accumulation of these important compounds in microbes is not yet possible. Although the activity of a P450 cyclase was inferred to catalyze the final committed step in glyceollin biosynthesis forty years ago, the enzyme in question has never been conclusively identified. This study reports, for the first time, the identification of three cytochrome P450 monooxygenase cyclases that catalyze the final steps of glyceollin biosynthesis. Utilizing *P. sojae*-soybean transcriptome data, along with genome mining tools and co-expression network analysis, we have identified 16 candidate glyceollin synthases (GmGS). Heterologous expression of these candidate genes in yeast, coupled with *in vitro* enzyme assays, enabled us to discover three enzymes capable of producing two glyceollin isomers. GmGS11A and GmGS11B catalyzed the conversion of glyceollidin to glyceollin I, whereas GmGS13A converted glyceocarpin to glyceollin III. The functionality of these candidates was further confirmed *in planta* through gene silencing and overexpression in soybean hairy roots. This groundbreaking study not only contributes to the understanding of glyceollin biosynthesis, but also demonstrates a new synthetic biology strategy that could potentially be scaled up to produce valuable molecules for crop and disease management.

## Introduction

Glyceollins are prenylated pterocarpans biosynthesized through the isoflavonoid branch of the phenylpropanoid pathway in soybean. Glyceollin isomers have attracted much attention in recent years due to their medicinal effects on humans and animals^1, 2, 3^. The pharmacological properties of glyceollins include antioxidant, antidiabetic, anticancer, anti-inflammatory, neuroprotective activities^4, 5^, and the ability to block estrogen receptors^2, 6, 7^. Furthermore, the recurrent use of antibiotics in animal production has led to the emergence of antibiotic resistance, which is causing a global challenge for public health and food systems. Therefore, there is an increasing demand for alternative antibiotics use in feed additives in livestock agriculture^8^. Owing to their broad-spectrum antimicrobial properties, glyceollins are being proposed to replace antibiotics in the swine industry to reduce antibiotic-resistant microorganisms in the food supply^9, 10, 11^.

Glyceollins act as phytoalexins that provide an intricate defense system against biotic and potentially abiotic stresses in soybean^12, 13, 14^. One of the major biotic challenges of soybeans is the highly infectious oomycete, *Phytophthora sojae,* that causes a devastating stem and root rot disease resulting in billions of dollars of yield loss annually worldwide^15^. Glyceollins play a pivotal role in inhibiting the growth and spread of *P. sojae* and many other microbial pathogens during soybean infection, constituting an essential component of the innate immune system^16, 17^. Even though the deployment of single resistance genes, commonly referred to as *Rps* (Resistance to *P. sojae*), effectively confers resistance against specific races of *P. sojae*, the long-term sustainability of *Rps* genes as a sole defense strategy has been compromised due to the emergence of new pathogen races. Upon infection by race 1 of *P. sojae,* a more rapid upregulation of glyceollin biosynthesis occurs in soybean genotypes encoding Rps1K^18, 19^. Reduction of glyceollin accumulation in *P. sojae* resistant soybean lines (*Rps1*) has been found to breakdown Rps-mediated resistance, demonstrating the importance of glyceollins to both race- and non-race-specific resistance^16, 20^.

The biosynthesis of glyceollin isomers involves at least 15 enzyme-catalyzed steps, with each enzyme encoded by multi-gene families^21^. They are produced via the legume-specific isoflavonoid branch of the phenylpropanoid pathway in soybean. The first step of glyceollin biosynthesis is the 2′-hydroxylation of the isoflavone aglycone, daidzein (Fig. 1). The successful demonstration of *de novo* biosynthesis of daidzein utilizing a yeast platform^22^ offers compelling evidence in favor of employing microbial systems for the synthesis of value-added bioactive molecules, including glyceollins. Most of the biosynthetic enzymes up to the penultimate enzymes G4DT^23^ and G2DT^24^ for the biosynthesis of glyceollin have been functionally characterized in soybean. Several of these isoflavonoid biosynthetic enzymes, belong to the cytochrome P450 monooxygenase (P450) enzyme families, including C4H, IFS, I2’H, and 3,9 DPO (Fig. 1). The enzyme involved in the final step in the biosynthesis of the major glyceollin isomers is suggested to be a P450 cyclase, GS^25^. Crude microsomal extracts were found to convert glyceollidin and glyceocarpin to their corresponding glyceollins through the cyclization of their prenyl residues, thereby completing the synthesis of this important class of phytoalexins. Despite the fact that these advancements had been made almost four decades ago, the enzyme and the precise cyclization mechanism involved in this process remained elusive.

**Fig. 1.**
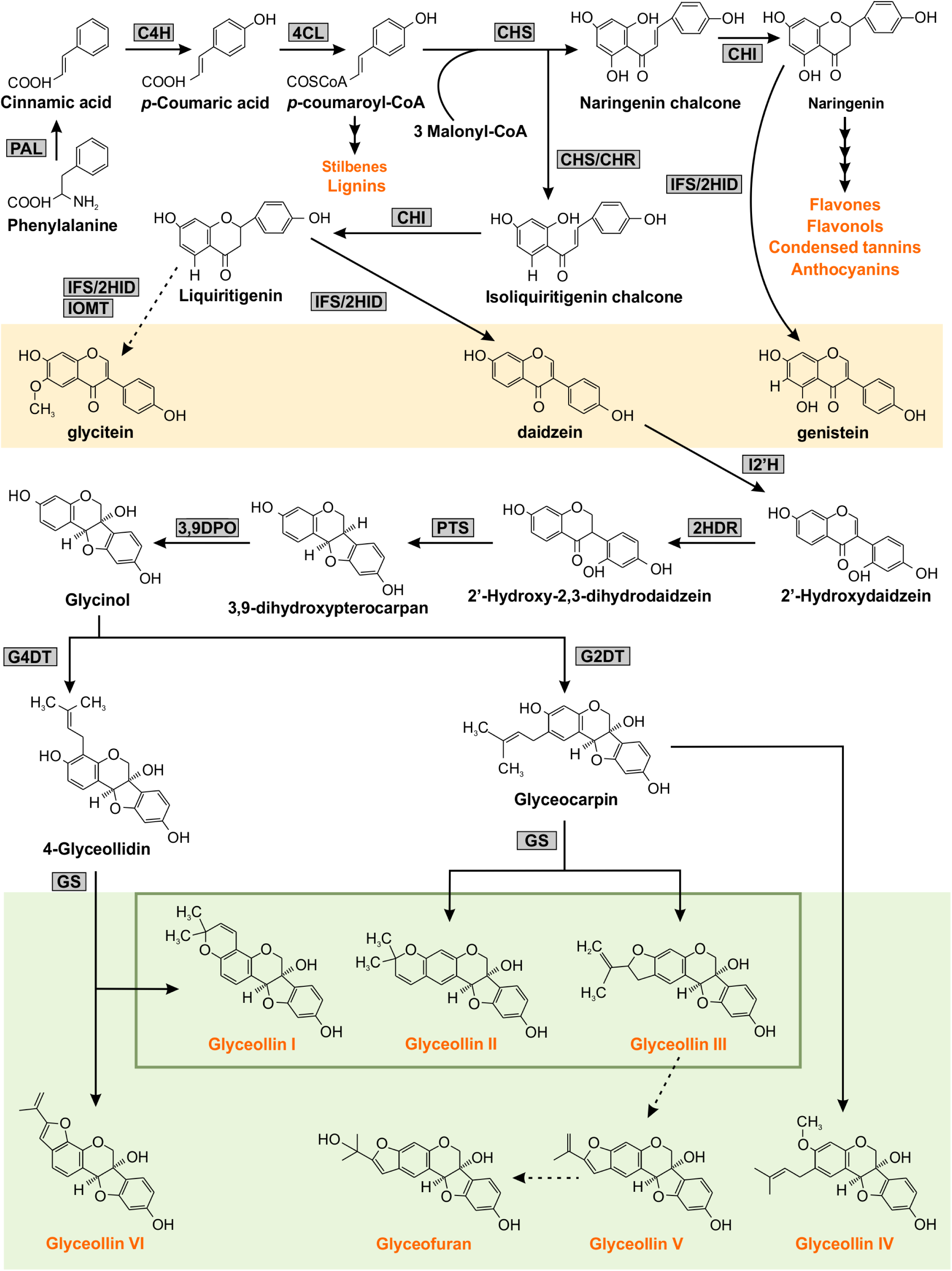
Glyceollin biosynthetic pathway in soybean. The multiple arrows indicate multiple steps and the dotted arrow indicates speculated steps in the pathway. The major metabolites synthesized from the phenylpropanoid pathway are shown in orange text. The three major isoflavone aglycones are highlighted in orange. PAL phenylalanine ammonia-lyase, C4H cinnamate-4-hydroxylate, 4CL 4-coumarate-CoA-ligase, CHS chalcone synthase, CHR chalcone reductase, CHI chalcone isomerase, IFS 2-hydroxyisoflavanone synthase, 2HID 2-hydroxyisoflavanone dehydratase, IOMT isoflavone Ο-methyltransferase, I2′H Isoflavone 2′-hydroxylase, 2HDR 2′-hydroxydaidzein reductase, PTS pterocarpan synthase, 3,9 DPO 3,9-dihydroxypterocarpan 6a-monooxygenase, G4DT Glycinol 4-dimethylallyltransferase, G2DT Glycinol 2-dimethylallyltransferase, GS glyceollin synthase. Glyceollin isomers are highlighted in green.

In this study, we provide a comprehensive characterization of three P450 enzymes classified as members of the CYP71 and CYP82 families, which exhibit co-expression with *GmIFS2* in response to *P. sojae* infection. We expressed these genes in a heterologous yeast system and investigated their potential enzymatic activity towards the substrates glyceocarpin and glyceollidin. We successfully identified *GmGS11A, GmGS11B* and *GmGS13A* as genes encoding the GSs. GmGS11A and GmGS11B convert 4-glyceollidin to glyceollin I, whereas GmGS13A converts glyceocarpin to glyceollin III. These conversions involve the formation of 2,2-dimethylchromen and 2-isopropenyldihydrofuran rings. This is the first report of a P450 with oxidative cyclization activity from the Fabaceae family. The discovery of the *GmGS* genes provides intriguing opportunities for the application of metabolic engineering strategies to manufacture value-added glyceollins in microbial systems.

## Results

### Identification of candidate GSs

Since glyceollin biosynthesis is an induced stress response, to identify the candidate *GmGS* genes, we searched transcriptomic datasets available in the public domain from five soybean-*P. sojae* interaction studies in which *P450* genes were upregulated during infection^16, 26, 27, 28, 29^. These datasets contained samples collected at time points ranging from 0.5 to 48 hours post *P. sojae*-infection, and included soybean cultivars that differ in resistance against stem and root rot diseases (Table S1). Analysis of differentially expressed genes (DEG) in each of these studies and comparing the DEG between the studies identified 21 common GmP450s that were upregulated in all five transcriptomic studies (Fig. 2). Filtering out the previously characterized genes such as *IFS* and *C4H* and potential *I2’H* genes, 16 GmGS candidates encoded by 15 genes were selected for further analysis (Table 1).

**Fig. 2.**
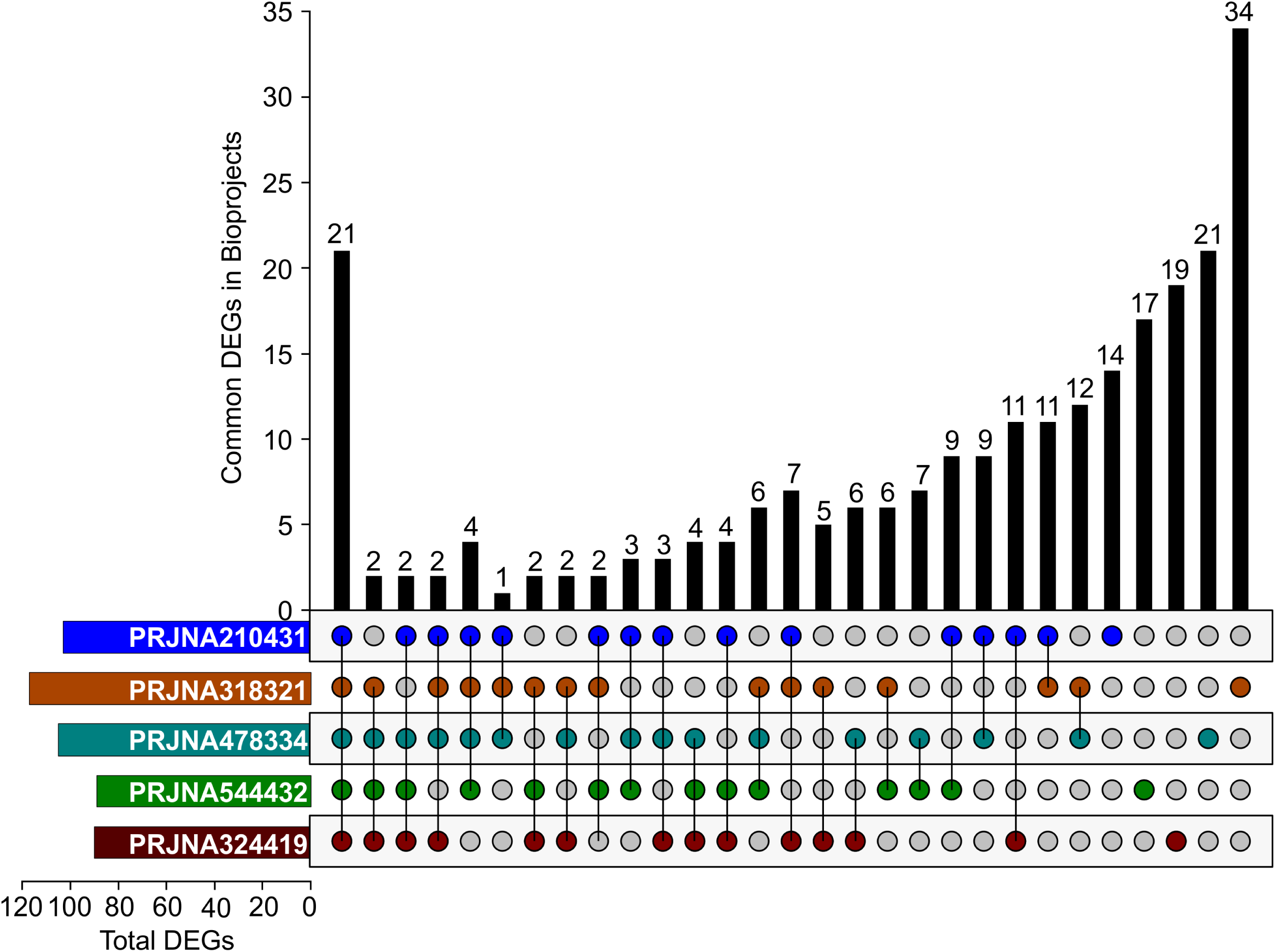
Gene expression analysis in soybean-*P. sojae* interaction. UpSet plot illustrating the expression patterns of GmP450 genes that are either common or unique across various bioprojects used in this investigation. Utilizing the publicly available RNAseq datasets involving soybean-*P. sojae* interaction (bioprojects: PRJNA324419, PRJNA544432, PRJNA210431, PRJNA318321, and PRJNA478334), differentially expressed *GmP450*s within each dataset were analyzed through pairwise comparisons. The diagram highlights distinct sets of differentially-expressed *GmP450*s post-infection with *P. sojae*, indicating the genes unique to a particular dataset or those shared among multiple datasets.

**Table 1:** Detailed information of *GmGS* candidates.

| Gene name | Gene locus | CYP name | Gene location | Predicted molecular Weight (kDa) | Coding DNA sequence (bp) | Splice variants | Number of introns |
| --- | --- | --- | --- | --- | --- | --- | --- |
| *GmGS01A* | *Glyma.01G135200* | CYP82A4 | Chr01: 46807741..46812089 | 58.8 | 1581 | 1 | 1 |
| *GmGS3B* | *Glyma.03G142100* | CYP93A3 | Chr03: 36982726..36984888 | 58.0 | 1533 | 1 | 1 |
| *GmGS05A* | *Glyma.05G147100* | CYP78A58 | Chr05: 34150611..34153212 | 57.3 | 1527 | 1 | 1 |
| *GmGS05B* | *Glyma.05G022100* | CYP75B43 | Chr05: 1932583..1936417 | 57.5 | 1536 | 1 | 2 |
| *GmGS08B-1* | *Glyma.08G140600* | CYP736A31 | Chr08: 10835711..10837975 | 56.9 | 1497 | 2 | 1 |
| *GmGS08C-1* | *Glyma.08G193900* | CYP90D12 | Chr08: 15702236..15707181 | 54.4 | 1425 | 2 | 7 |
| *GmGS10A* | *Glyma.10G203500* | CYP76F18 | Chr10: 43557513..43559769 | 58.4 | 1572 | 1 | 1 |
| *GmGS11A* | *Glyma.11G062500* | CYP71D145 | Chr11: 4723140..4726670 | 57.4 | 1518 | 1 | 1 |
| *GmGS11B* | *Glyma.11G062600* | CYP71D8 | Chr11: 4730070..4732519 | 57.6 | 1515 | 2 | 1 |
| *GmGS13A* | *Glyma.13G285300* | CYP82A2 | Chr13: 37999351..38002170 | 58.7 | 1569 | 1 | 1 |
| *GmGS13B* | *Glyma.13G068800* | CYP82A3 | Chr13: 15920188..15925586 | 59.8 | 1584 | 3 | 2 |
| *GmGS16A* | *Glyma.16G195600* | CYP71AU9 | Chr16: 35935519..35938084 | 58.0 | 1554 | 1 | 1 |
| *GmGS19B* | *Glyma.19G144700* | CYP93A2 | Chr19: 41024728..41027051 | 57.2 | 1509 | 1 | 1 |
| *GmGS19C* | *Glyma.19G146800* | CYP93A41 | Chr19: 41200476..41202854 | 57.9 | 1530 | 1 | 2 |
| *GmGS20A-1* | *Glyma.20G159700* | CYP715A12 | Chr20: 39786056..39789417 | 58.8 | 1560 | 3 | 3 |
| *GmGS20A-3* | *Glyma.20G159700* | CYP715A12 | Chr20: 39786056..39789417 | 43.0 | 1146 |  | 2 |

### Collinearity analysis of *GmGSs* with other legumes

The Fabaceae family evolved approximately 60-70 million years ago^30^. To detect syntenic and collinear relationship of soybean with its wild relative and other legumes, we constructed collinearity syntenic maps of soybean with barrel clover (*Medicago truncatula),* pea (*Pisum sativum),* common bean (*Phaseolus vulgaris),* lentil (*Lens culinaris),* chick pea (*Cicer arietinum)* and wild soybean (*Glycine soja*). The candidate *GmGS* gene pairs within the significant collinear blocks were identified (Fig. 3, Table S2). Our analysis identified 13 and six colinear gene pairs in *G. max* - *G. soja* and *G. max* – *P. vulgaris*, respectively. Four soybean *GmGS* genes paired in both barrel clover and chickpea collinear blocks, while two paired in each pea and lentil. As expected, a higher number of *GmGS* gene pairs was observed in *G. max* and *G. soja,* the only other legume species capable of producing glyceollins^31^. Among other legumes, larger number of gene pairs in soybean and common bean suggests a closer evolutionary relationship between these two species. Furthermore, the conserved collinear pairs, especially involving *GmGS08C-1* and *GmGS11A* across all legume species, provide insight into the evolutionarily diverse legume species.

**Fig. 3.**
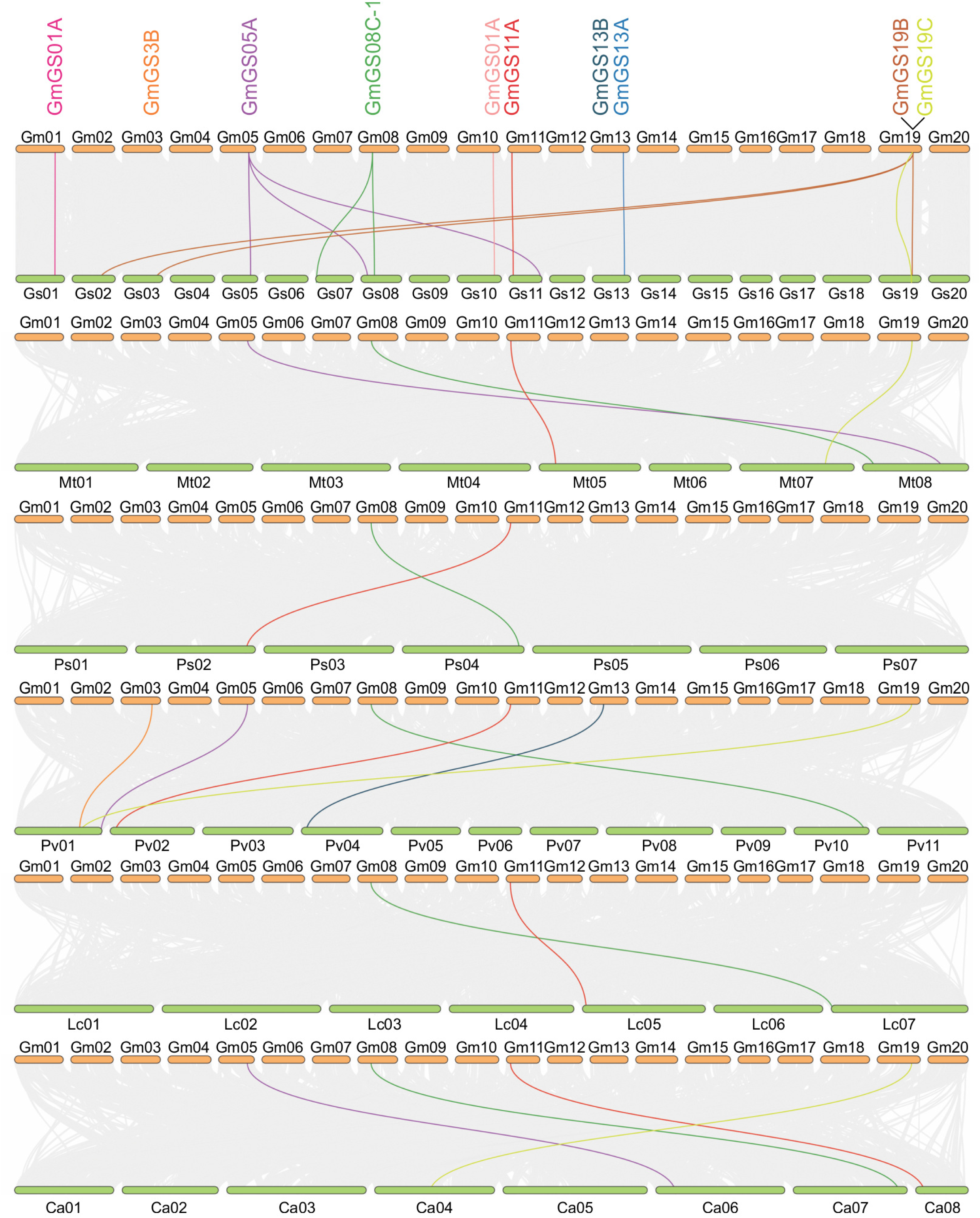
Synteny analyses of candidate *GmGS* and genes from other legume species. The collinearity blocks within soybean and other specie genomes are shown with gray lines. The syntenic *GmGS* gene pairs between soybean and other legume species are indicated with colored lines. Gm *Glycine max*, Gs *Glycine soja*, Ps *Pisum sativum*, Pv *Phaseolus vulgaris*, Lc *Linum culnaris*, Mt *Medicago truncatula*, Ca *Cicer arietinum*

### Gene co-expression network analysis reduces 15 *GmGS* candidates to three

Co-expression network analysis represents a powerful tool for prioritizing genes that exhibit similar temporal and spatial expression patterns, suggesting related functions such as involvement in common metabolic pathways. The biosynthesis of glyceollin requires of at least 15 enzymatic steps, and each enzyme contains multiple isoforms. A search for genes encoding all the enzymes and their isoforms identified 71 genes that possibly lead to glyceollin biosynthesis (Table S3). A co-expression analysis of 15 candidate *GmGS* genes and the 71 glyceollin biosynthetic genes was conducted. Since *GmGSs* are predicted to be P450s^25^, we also included all the *GmP450s* in the co-expression study. The Soybean Expression Atlas v2 that contains 10 transcriptomic bioProjects focused on soybean-*P. sojae* interactions was utilized for the expression analysis^32^. The analysis produced a co-expression score based on correlation between each pair of genes in the samples (Table S3). A total of 118 genes exhibiting significant coefficients (r ≥ 0.6) were included in creating the network. The results revealed two major and several minor networks (Fig. 4; Supplementary Fig. S1). The main network contained six *GmGS* gene candidates (*GmGS01A, GmGS11A, GmGS11B, GmGS13A, GmGS16A* and *GmGS19C*), where only three genes formed a tight network and showed significant correlations with multiple glyceollin biosynthetic genes. *GmGS11A* expression correlated with the expression of *GmCHS1, GmCHS10, G4DT, GmGS11B, GmGS13A, GmGS19C, GmIFS2, GmCHR14,* and *Glyma.15G156100* (CYP81). *GmGS11B* displayed positive correlations with *GmGS11A, GmCHS1, GmGS13A, GmIFS2, G2DT, G4DT, GmPTS1, GmGS16A, GmIFR,* and *Glyma.15G156100*. *GmGS13A* expression was positively correlated with *GmGS11A, GmGS11B, GmCHS1, GmCHS10, GmCHS5, GmIFR, G2DT, G4DT, GmPTS1, GmIFS2,* and *Glyma.15G156100*. The expression of three candidate *GmGS*s, *GmGS11A, GmGS11B* and *GmGS13A,* show significant correlation with *GmIFS2* (r = 0.6389 to 0.7607) and the penultimate genes *G4DT* and *G2DT* (r = 0.6119 to 0.6623) (Table S2). Additionally, *GmGS16A* and *GmGS01A* showed correlations with each other and with *GmCHI1A* and *GmIFR*. Even though *GmGS3B, GmGS19C* and *GmGS20A* were the part of the network, their expression was correlated with only one or two genes. This refined analysis emphasizes the specific associations of *GmGS11A, GmGS11B,* and *GmGS13A* within the major co-expression network, thus narrowing the candidates to three genes.

**Fig. 4.**
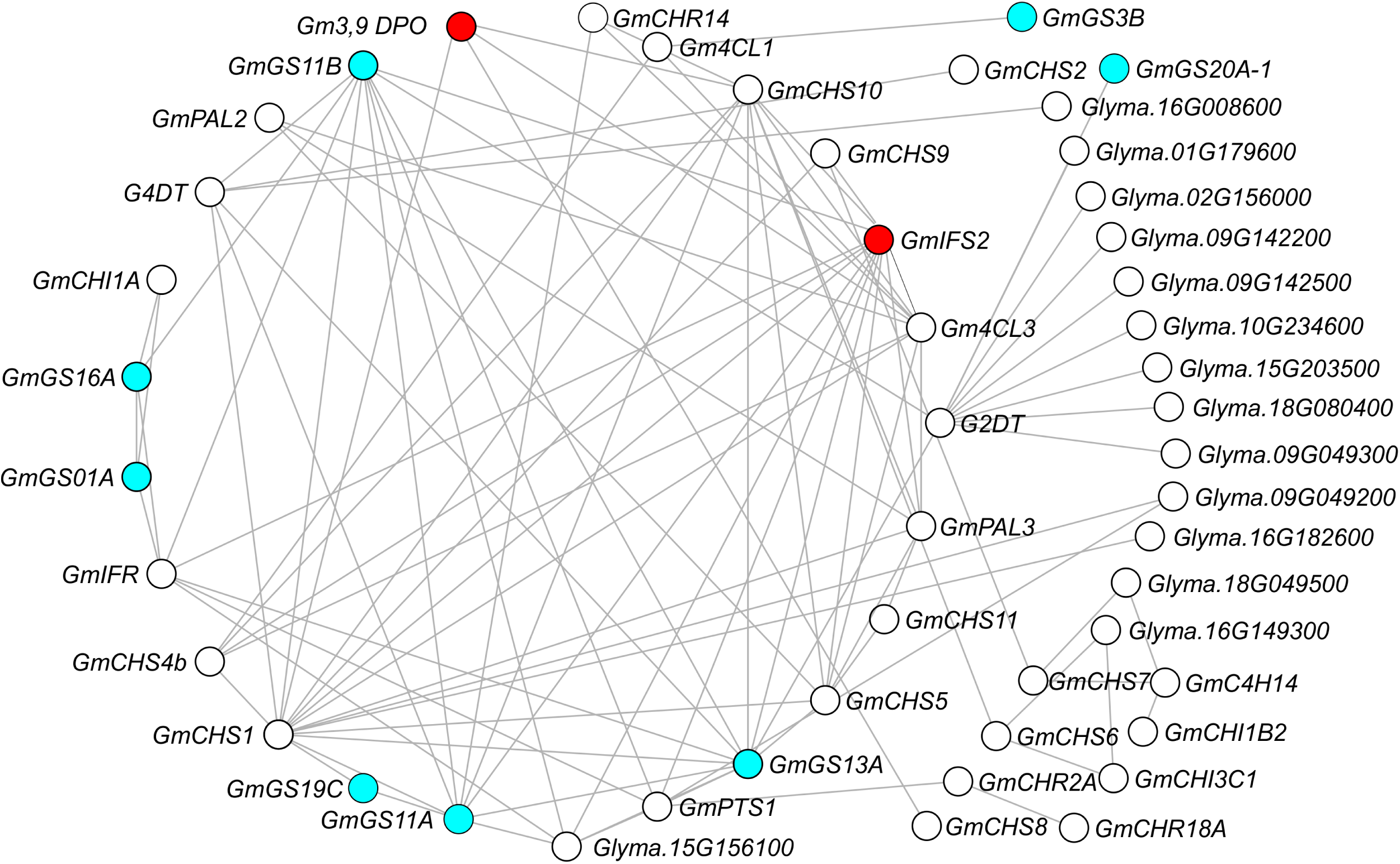
Co-expression network analysis of *GmGS* candidates with glyceollin biosynthetic genes and *GmP450s*. Red circles indicate *IFS2* and *3,9-dihydroxypterocarpan 6A-monooxygenase* and blue circles show *GmGS* candidates.

GmGS11A and GmGS11B are encoded by *Glyma.11G062500* and *Glyma.11G062600,* respectively that reside tandemly on chromosome 11 in soybean. On the other hand, GmGS13A encoded by *Glyma.13G285300* is located on chromosome 13. Each of the three *GmGS*s contain a single intron in their genomic sequence. Despite the fact that *GmGS11A* and *GmGS11B* display very high (94%) sequence identity at the nucleotide level, there exists a significant difference in the size of their introns (1762 nt for *GmGS11A* and 627 nt for *GmGS11B*).

### Identification of three GmGSs with glyceollin synthase activity using *in vitro* yeast assay

The final committed step of glyceollin biosynthesis includes oxidation and cyclization reactions, converting glyceollidin and glyceocarpin into glyceollin I and glyceollin II/III, respectively (Figs. 1 and 5a, b). To investigate the biochemical activity of the candidate GmGSs, we used previously purified glyceocarpin^24^. To produce glyceollidin, glycinol was used as a substrate for G4DT^23^. Production of glyceollidin was confirmed prior to the GmGS enzyme assay. Microsomes prepared from yeast cells co-expressing candidate GmGS and *Lotus japonicus* cytochrome P450 reductase 1 (LjCPR1) were used *in vitro* for measuring substrate depletion and new compound formation, as this method has proven to be successful previously^33^. Microsomal protein preparation from yeast cells expressing only LjCPR1 (empty vector) and reactions with no enzyme (substrate only) were used as controls (Fig. 5c-d and k-l). A complete depletion of glyceollidin and formation of glyceollin I was detected when microsomes from yeast cells expressing GmGS11A and GmGS11B were used (Fig. 5e, f). The major enzymatic product had the same retention time analyzed by high resolution mass spectrometry with a molecular formula of C_20_H_18_O_5_ (*m/z* 321.1121, [M-H_2_O+H]^+^) compared to a synthetic authentic standard of glyceollin I (Fig. 5i). Similarly, when glyceocarpin was used as a substrate, a complete depletion of glyceocarpin and the formation of a new product was detected only in the reaction containing GmGS13A (Fig. 5o). The observed product in this enzymatic reaction is the first of three isomers to elute by liquid chromatography (Fig 5p) that contains the molecular formula of C_20_H_18_O_5_ (*m/z* 321.1121, [M-H_2_O+H]^+^), and possesses characteristic product ion spectra (Fig 5r) that matched previous reports for glyceollin III^34^. This indicates that GmGS13A catalyzes the conversion of glyceocarpin to glyceollin III. In all three reactions, only a single product was detected, suggesting a direct cyclization of the substrate to the final products. To determine if other candidate GmGSs could use glyceocarpin or glyceollidin as a substrate, we performed enzyme assays with the 13 remaining candidate proteins (Table 1). None of them were able to convert either glyceocarpin or glyceollidin to their corresponding glyceollins.

**Fig. 5.**
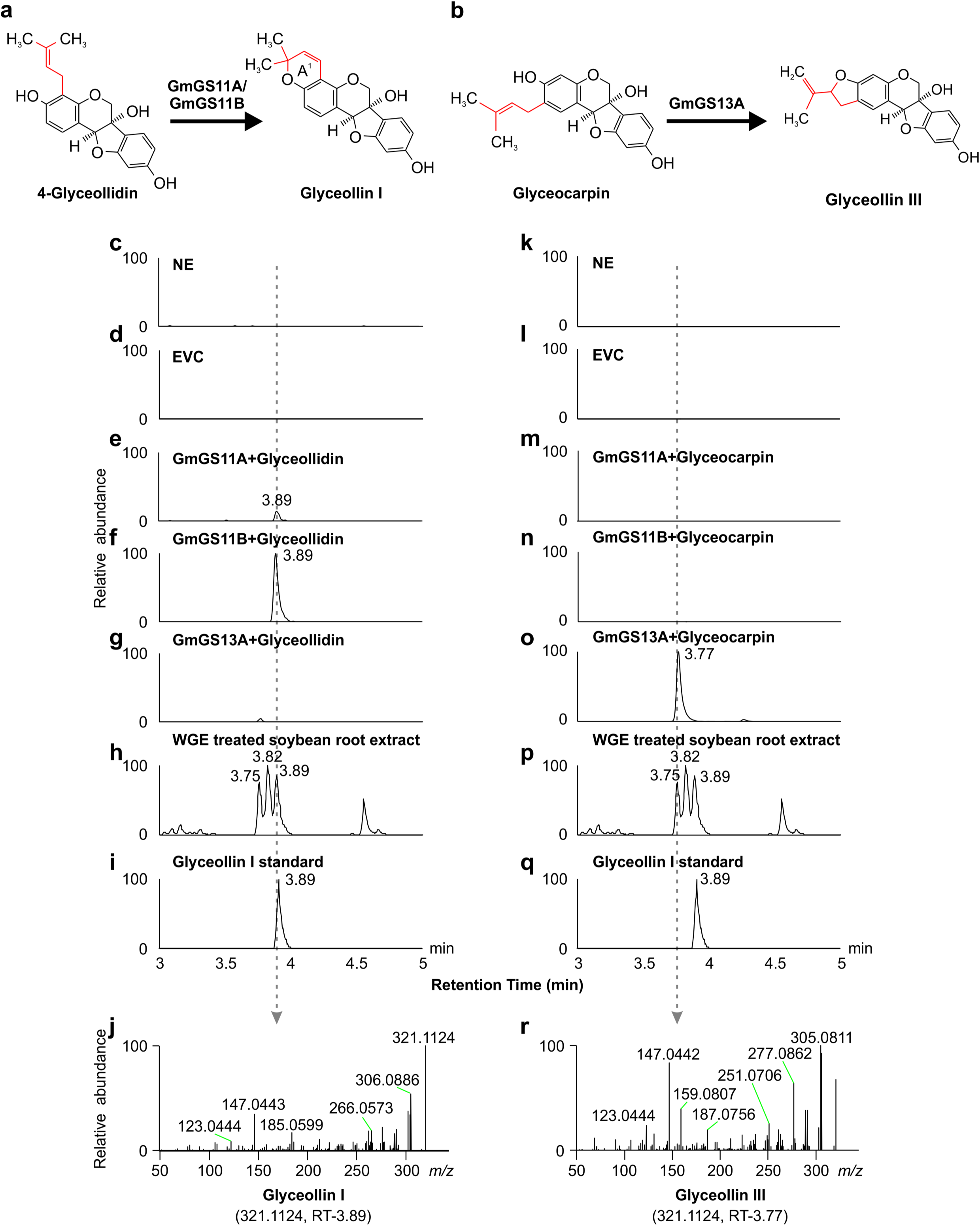
*In vitro* enzyme assays for glyceollin synthase activity. Microsomes were isolated from *S. cerevisiae* strain BY4742 expressing *LjCPR1* (empty vector control) or *LjCPR1* and *GmGS* followed by enzyme assay and LC-MS analysis. Glyceollins were monitored in samples as the [M-H2O+H]+ ion at *m/z* 321.1121 (±3 ppm) with the isomers being only distinguishable by retention time. **a.** Enzymatic reaction showing conversion of glyceollidin to glyceollin I. **b.** Enzymatic reaction showing conversion of glyceocarpin to glyceollin III. Extracted ion chromatogram (EIC) and product ion spectra for glyceollin isomers (*m/z* = 321.1121) are shown. **c, k.** No enzyme control (NE), **d, l.** Empty vector control (EVC), **e.** Microsomes containing recombinant GmGS11A+glyceollidin substrate, **f.**Microsomes containing recombinant GmGS11B+glyceollidin substrate, **g.**Microsomes containing recombinant GmGS13A+glyceollidin substrate, **h**, **p.** WGE-treated soybean root extracts, **i, q.** Glyceollin I standard, **j.** Product ion spectrum for glyceollin I (retention time = 3.89). **m.** Microsomes containing recombinant GmGS11A+glyceocarpin substrate, **n.** Microsomes containing recombinant GmGS11B+glyceocarpin substrate, **o.** Microsomes containing recombinant GmGS13A+ glyceocarpin substrate, **r.** Product ion spectrum for the peak eluting earlier (retention time = 3.75 min) than glyceollin I.

### *GmGS11A, GmGS11B* and *GmGS13A* independently alter glyceollin level *in planta*

To confirm whether the *GmGS* genes function in glyceollin biosynthesis, we performed over-expression (OE) and RNAi silencing (Si) using a soybean hairy root system. The coding sequence of the genes for OE, and gene-specific fragments for Si, were selected and cloned into the respective vectors. Due to the high nucleotide sequence identity between *GmGS11A* and *GmGS11B* (94%), an Si fragment could not be designed to target the two genes independently. Both RNAi and OE vectors consist of a separate cassette that express the eGFP marker. Transgenic hairy roots were selected on the basis of eGFP fluorescence and were maintained individually as independent transgenic lines (Fig. 6a, b). Each transgenic hairy root line was treated with AgNO_3_, a chemical elicitor that mimics *P. sojae* infection. Silencing and overexpression of the target genes in multiple independent transgenic hairy roots (AgNO_3_-treated and control) were assessed by comparing their transcript levels. The effect of AgNO_3_ treatment on glyceollin accumulation in the control, Si and OE lines were also monitored. The results revealed that GmGS11A-Si roots (n=4 biological replicates) contained significantly reduced levels of *GmGS11A* and *GmGS11B* transcripts compared to the controls (Fig. 6c). Silencing of *GmGS11A* and *GmGS11B* significantly reduced the amounts of glyceollin I in GmGS11A/11B-Si roots compared to the controls, while glyceollin II and glyceollin III levels were unaltered (Fig. 6d). The GmGS13A-Si lines (n=4 biological replicates) displayed significantly reduced levels of *GmGS13A* transcripts as well as glyceollin III levels (Fig. 6e, f). No change in the levels of glyceollin I and glyceollin II were detected in GmGS13A-Si roots. The overexpression of *GmGS11A, GmGS11B*, and *GmGS13A* led to increased level of transcripts in the respective transgenic hairy roots compared to the control (Fig. 6g-i). While no change in the accumulation of glyceollin isomers was observed in GmGS11A-OE roots, GmGS11B-OE and GmGS13A-OE roots showed significantly higher accumulation of glyceollin I and glyceollin III, respectively (Fig.6j-l). Additionally, GmGS11B-OE accumulated significantly higher levels of glyceollin II (Fig. 6k). These results verify our *in vitro* enzyme assay results that GmGS11B and GmGS13A are involved in glyceollin I and glyceollin III biosynthesis, respectively.

**Fig. 6.**
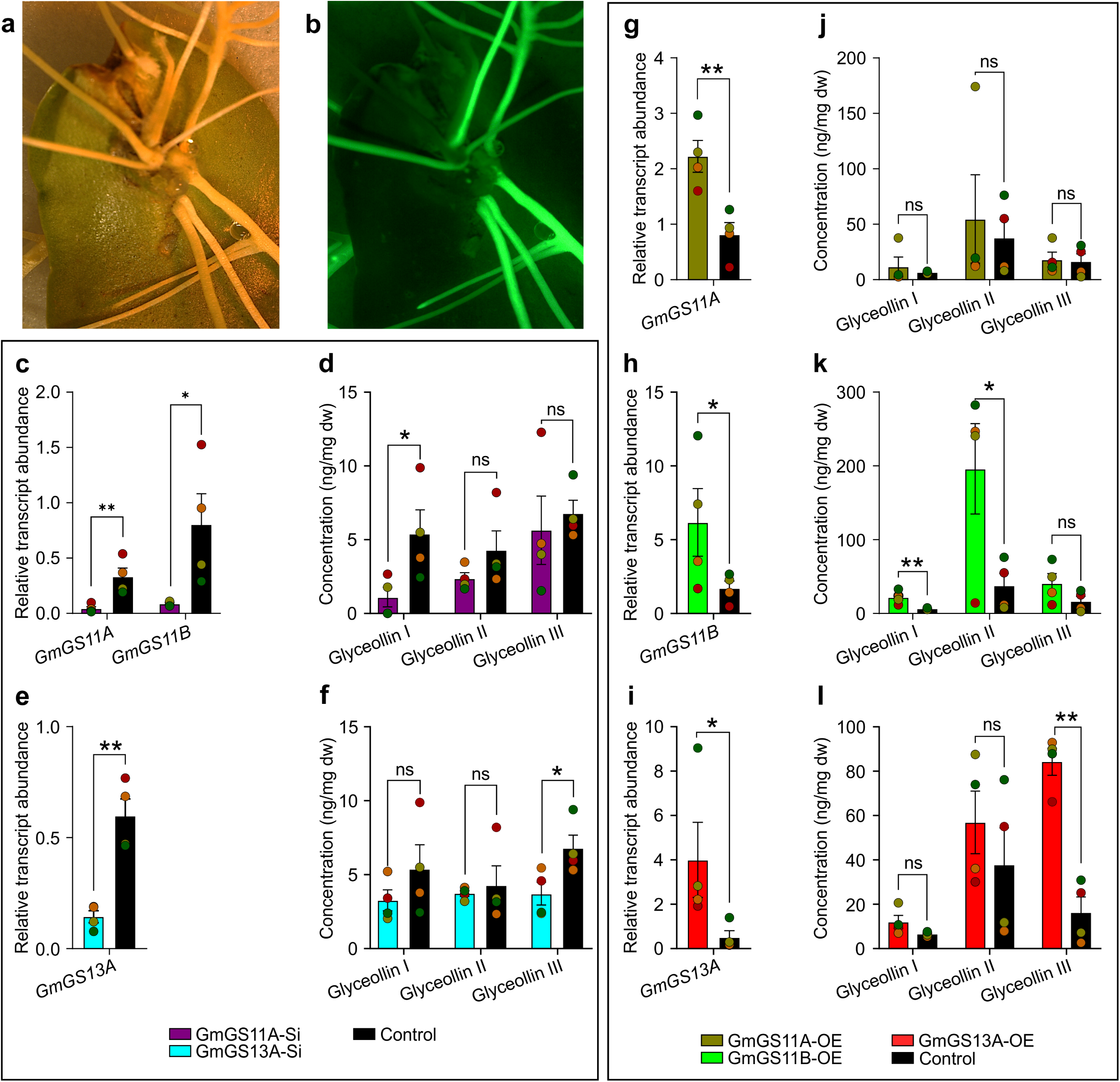
Functional characterization of *GmGS11A*, *GmGS11B* and *GmGS13A in planta*. **a.** Soybean hairy roots transformed with RNAi construct photographed under white light. **b.** The image shown in (a) under UV light with a GFP filter. Roots showing green fluorescence were harvested and cultured separately as independent transgenic lines. **c.** *GmGS11A* and *GmGS11B* transcript levels in silenced (Si) roots compared to control hairy roots analyzed using quantitative RT-PCR. **d.** Glyceollin levels in GmGS11A-Si and GmGS11B-Si lines compared to control hairy roots. **e.** *GmGS13A* transcript levels in GmGS113A-Si roots compared to control hairy roots. **f.** Glyceollin levels in GmGS13A-Si lines compared to control hairy roots. **g, h, i.** Relative transcript abundance of *GmGS11A, GmGS11B and GmGS13A* in OE transgenic lines. **j, k, l.** Glyceollin levels in OE lines compared to control hairy roots. Relative transcript abundance and glyceollin concentration indicate mean values of four biological replicates ± SEM. Asterisks indicate the statistical significance of *t*-test. * *P <* 0.05, ** *P <* 0.01, ns not significant (*P >* 0.05).

## Discussion

In addition to their significance in soybean agriculture in protecting against microbial pathogens such as *P. sojae*, glyceollins have drawn much attention as prospective therapeutics and antibiotics for humans and animals due to their pharmacological activities^9, 11, 35^. In healthy soybean plants, they are found in trace amounts; but when stressed, their biosynthesis is upregulated^25^. Currently, mixtures of glyceollin are generally extracted from soybeans challenged with the food-safe fungus *Aspergillus sojae* or produced from biosynthetic intermediates by semi-synthesis^36, 37, 38^. A complex eight-step total chemical synthesis of glyceollins I and II has been reported recently (Ciesielski and Metz, 2020). While engineering their biosynthesis into microorganisms could provide a ‘greener’ and more economical production method, the genes responsible for catalyzing the last committed step of each of their biosynthetic pathways have remained undiscovered. Here, we have identified three *GmGS* genes and characterized the activities of their encoded enzymes, which have long been speculated but never demonstrated.

Oxidative cyclization reactions are reported to be catalyzed by Diels-Alderases, however, they require flavin adenine dinucleotide^39^. The NADPH- and oxygen-dependent conversion of the prenylated pterocarpans to glyceollin isomers suggested that a P450 is involved^25^. By mining the publicly available five transcriptomic datasets of the soybean-*P. sojae* interaction (Table S1), we identified 21 P450 genes that were induced upon *P. sojae* infection (Fig. 2). As glyceollin production is a stress-induced response, the repetitive appearance of a set of the same genes in different soybean cultivars under a variety of experimental conditions provided confidence to proceed with these gene candidates. Based on the results of the co-expression analysis with the known isoflavonoid genes, we selected a handful of genes for the detailed functional analysis (Table1). Enzymatic assays using yeast microsomes expressing the candidate GmGSs with both glyceocarpin and glyceollidin as substrates identified GmGS11A and GmGS11B. These two enzymes specifically converted glyceollidin to glyceollin I, while GmGS13A converted glyceocarpin into glyceollin III (Fig. 5).

Endogenous glyceollin I, glyceollin II, and glyceollin III are the three major glyceollins produced by soybean roots challenged with wall glucan elicitor. These glyceollin isomers have previously been shown to elute by HPLC on a C18 column in reverse order^16, 34, 35, 40, 41, 42^. Accordingly, our analysis of WGE-treated soybean roots (Fig. 5h, p) show three isobaric, yet chromatographically resolved peaks for compounds of formula C_20_H_18_O_5_, with the later eluting peak confirmed as glyceollin I through comparison to the authentic synthetic standard (Fig. 5i, q). In the case of GmGS11A and GmGS11B, the reaction product was identified as glyceollin I by comparison of its LC retention time, chemical bared formula determined by accurate mass, and high resolution product ion spectra to those of the authentic synthetic standard. For GmGS13A, although there is no authentic standard of glyceollin III available, three lines of evidence strongly suggest that the identity of the reaction product is glyceollin III: i) the chemical formula determined by accurate mass, ii) the product ion spectra with high similarity to glyceollin I (Supplementary Fig. S2), and iii) the chromatographic retention time relative to glyceollin I matching the earliest eluting peak in WGE treated samples. The substrate and the product of GmGS13A, have the molecular formulas C_20_H_20_O_5_ and C_20_H_18_O_5,_ respectively, suggesting a cyclization of the prenyl chain to the proximal alcohol group on the A ring (Fig. 5b). Cyclization of the prenyl chain to the alcohol group of the A-ring system of glyceollidin leads to the formation of the new A^1^ ring of glyceollin I. The difference between the substrates glyceollidin and glyceocarpin is the position of the prenyl chain on the A ring, and thus an equivalent cyclization between the prenyl chain and the alcohol group of glyceocarpin’s A ring system would generate glyceollin III ^34, 40^.

Both GmGS11A and GmGS11B belong to the CYP71 family, while GmGS13A is a CYP82. The dual catalytic activities of two CYP71D family members involved in tanshinone synthesis in *Salvia miltiorrhiza*^43^ and an elicitor-inducible CYP71D20 in *Nicotiana benthamiana* have been reported^44^. CYP71D20 catalyzes two hydroxylation reactions of its substrate 5-*epi*-aristolochene. Oxidative scaffold formation and cyclization of alkaloids have also been reported^45^. More recently, a new dual function of GmCYP82D26 has been reported, which seems to have a shared role with IFS, CYP93C2^46^. Nevertheless, the cyclase role of plant P450 is not only rare but rather exceptional. Prenyl cyclization on rotenoids, pterocarpans and coumarins have been reported earlier^47^. Although an oxidative cyclization of prenylated rotenoid, rot-2′-enomic acid, to deguelin is catalyzed by a deguelin cyclase in *Tephrosia vogelli,* this deguelin cyclase is not a P450 and that the reaction does not require a cofactor^48^.

The *in planta* role of GmGS11A, GmGS11B and GmGS13A in glyceollin biosynthesis, was demonstrated by overexpressing and silencing these genes in soybean hairy roots and monitoring the effect on glyceollin accumulation upon elicitor treatment. Simultaneously silencing *GmGS11A* and *GmGS11B* reduced their transcripts and glyceollin I metabolites in multiple independent transgenic lines, while the amounts of glyceollin II and glyceollin III remained unaltered (Fig. 6c, d). Likewise, the reduction and increase in *GmGS13A* transcripts in GmGS13A-Si and GmGS13A-OE roots significantly reduced or increased glyceollin III amounts, respectively, but had no significant effect on glyceollin I and glyceollin II (Fig. 6e, f, i, l). This finding provided evidence for the *in vivo* role of GmGS13A in glyceollin III biosynthesis. As expected, the overexpression of *GmGS11B* significantly increased the amounts of glyceollin I (Fig. 6h, k). However, the increased level of glyceollin II in GmGS11B-OE roots was surprising. GmGS11B was neither able to accept glyceocarpin as a substrate in the *in vitro* enzyme assay nor produce glyceollin III from glyceollidin (Fig 5f, n). Based on these results, we speculate that plant-specific factor such as physically associated protein(s)^49^ possibly alter the regiospecificity of GmGS11B to produce glyceollin II. Furthermore, an increase in transcript levels of *GmGS11A* was not accompanied by the higher accumulation of glyceollin I metabolites (Fig. 6g, j). It is possible that glyceollin I produced by *GmGS11A* serves as an intermediate for the biosynthesis of another specialized metabolite in soybean. Previous pulse-chase experiments supported that glyceollin I, like several other phytoalexins, is not a pathway ‘end product’, rather it is converted into one or more unknown molecules, and this turnover and glyceollin biosynthesis are accelerated by elicitation^50^.

The identification of exclusive collinearity pairs in soybean and its wild counterpart *G. soja*, highlights their close genetic relationship. This genomic connectivity between soybean and *G. soja* suggests the importance of the evolution and maintenance of phytoalexin biosynthesis mechanisms, reflecting shared adaptive strategies in response to challenges by common pathogens. The varying syntenic patterns with other legumes further align with the lineage-specific phytoalexins produced by different legumes, highlighting the complex evolutionary dynamics within the Fabaceae family.

In conclusion, this discovery identified the long-speculated P450s that catalyze the ultimate steps of the glyceollin biosynthetic pathway. It also informs on the catalytic diversity of the plant P450 family and provides new possibilities in biomanufacturing of glyceollin with medicinal value in microbial or plant systems. Equally important, the above mentioned findings should greatly accelerate the identification of new molecular markers for resistance breeding in soybean.

## Methods

### Plant growth conditions and treatments

Soybean (*G. max* L. Merr) cv. W82 seeds were obtained from a collection of Agriculture and Agri-Food Canada (London Research and Development Centre). Seedlings were grown in water-saturated vermiculite supplemented with 0.3 g/L of fertilizer (20-15-20) in a growth chamber with a 24 h dark cycle at 22°C and 60% relative humidity. Seven-day-old etiolated seedlings were either inoculated with 1.0 mM AgNO_3_ or infected with actively-growing 5-day-old *P. sojae* (race 25) mycelia cored out from the leading edge of the culture grown on a V8-agar medium. The hypocotyl tissue was harvested 24 hours following the treatment, flash-frozen in liquid nitrogen and then stored at −80°C until used.

### RNA extraction, RT-PCR and quantitative RT-PCR

Total RNA was isolated from *P. sojae* infected tissues using RNeasy Plant Mini Kit (Qiagen, USA) with on-column RQ1 RNase-Free DNase 1 treatment (Promega, USA). cDNA was synthesized using the SuperScript^TM^ IV First-Strand Synthesis System (ThermoFisher, USA) and used as a template for the amplification of full length *GmGS* candidates with gene-specific primers (Table S4).

### Plasmid construction

For gene cloning, all recombinant plasmids were constructed using Gateway technology (Invitrogen, USA). Genes of interest (GOI) were amplified using high-fidelity Platinum SuperFi DNA Polymerase (ThermoFisher, USA) with gene-specific *attB*-adapter primers compatible with the Gateway Technology (Table S4). Entry clones were obtained via a BP recombination reaction between the entry vector pDONR/Zeo and the *attB* PCR products. Following sequence verification, expression clones were assembled in an LR recombination reaction between the entry clones and destination vectors. For heterologous expression in yeast, GOIs were cloned into pESC-Leu2d-LjCPR1-GW (Khatri et al, 2023) and transferred to *Saccharomyces cerevisiae* strain BY4742. For RNAi and overexpression of GOIs, destination vectors pK7GWIWG2D (II),0 and pK7WG2D, respectively, were used. Recombinant RNAi and overexpression plasmids were transformed into *Agrobacterium rhizogenes* K599 by electroporation.

### Hairy root generation

Soybean hairy root generation was performed according to Subramanian et al (2005). Soybean cotyledons were inoculated with *Agrobacterium rhizogenes* K599 harboring gene of interest in either the overexpression or RNAi vector. Transgenic roots were selected 3-4 weeks post inoculation, under UV-illuminated NiKon SMZ25 stereo microscope (Nikon Instruments Inc, USA). Hairy roots were sub-cultured on root growth medium (4.33 g/L MS salts mixture, 3% sucrose, 0.2% gelzan (w/v) [pH 5.8], 2.5× MS vitamin mixture and 500 μg/mL of Timentin (GoldBio, USA)) until harvested for analysis.

### Heterologous expression and enzyme assay

Expression of recombinant GmGS proteins in yeast and microsome preparation was performed as described in Khatri et al (2023). For GmGS enzyme assay, microsomal proteins (1 mg) were incubated with 50 mM potassium phosphate buffer [pH 7.6] containing 1 mM NADPH and 1-5 μg of the substrate (glyceocarpin or glyceollidin) in a total volume of 200 µL. The mixture was incubated at 25°C for 6 hours with shaking. The reaction was stopped with an equal volume of chilled methanol and mixing vigorously for 10-15 seconds. The insoluble material was removed by centrifugation twice at 13,000*g*, 4°C for 5 min and the supernatant was subjected to LC-high resolution MS analysis.

### Metabolite extraction and LC-high resolution mass spectrometry analysis

Total isoflavonoid extraction from soybean hairy roots were performed as previously described^51^. A Q-Exactive Quadrupole Orbitrap mass spectrometer (Thermo Fisher Scientific, USA) coupled to an Agilent 1290 HPLC was used for LC-HRMS analysis. The HPLC system used a Accucore™ Phenyl-Hexylcolumn (2.1 x 50 mm, 2.6 μm) maintained at 35°C. Samples (2 μL each) were injected and run using a flow rate of 0.7 mL min^-1^. Mobile phase A consisted of water with 10 mM ammonium formate and 0.1% formic acid. Mobile phase B consisted of 5% water, 95% methanol, 5 mM ammonium formate and 0.1% formic acid. Mobile phase B was held at 2% for 0.75 min and increased to 35% over 0.5 min. B was increased to 65% over 2.75 min and finally to 100% over 3.5 min. It was maintained at 100% for 2.5 minutes before returning to 0% over 30 seconds and held for 1.5 min. Heated electrospray ionization (HESI) conditions used are as follows; spray voltage, 4.25 kV (ESI); capillary temperature, 400°C; probe heater temperature, 450°C; sheath gas and auxiliary gas 30 and 8 arbitrary units, respectively; S-Lens RF level, 45. The acquisition method consisted of both a full MS scan and parallel reaction monitoring (PRM) in positive ionization mod. The full MS scan monitored the mass range of *m/z* 100 to 1000, at 35,000 resolution, automatic gain control (AGC) target of 5×10^5^ and maximum injection time of 128 ms. The PRM scans used to collect product ion spectra of glyceollins I, II and III were acquired at a resolution of 17,500, AGC of 5×10^6^, maximum injection time of 64 ms. isolation window of 1.2 *m/z* at a normalized collision energies (NCE) of 50. Data analysis was performed using the manufacturer’s Xcalibur™ software (Thermo Fisher Scientific, USA) with a 1/x weighting on the calibration curve. The glyceollin I standard was purchased from Dr. Paul Erhardt, Department of Medicinal and Biological Chemistry, University of Toledo. Glyceollin I was used as a surrogate standard for the quantification of glyceollin II and III by monitoring all three as the [M-H2O+H]+ ion.

### In silico analysis

The co-expression network was constructed with all soybean P450s^52^, previously reported glyceollin biosynthetic genes (Table S2), along with *GmGS* candidates serving as bait. The network was generated using the CoExpNetViz tool with the default cutoff parameters^53^. Normalized expression data for GmP450s and other pathway genes from 10 transcriptomic studies on the *P. sojae*-soybean interaction (Table S2) were obtained from the Soybean Expression Atlas v2 (https://soyatlas.venanciogroup.uenf.br/)^32^. This data was utilized to generate the network. To identify correlations among all putative GmP450 genes, pathway genes and GmGS candidates, the analysis employed the Pearson correlation (r) method, with a correlation threshold set at the lower percentile rank of 5 and upper percentile rank of 95. Visualization of the co-expression was achieved using Cytoscape V3.10.1 (Smoot et al., 2011).

To conduct synteny analysis, we collected the genome and GFF3 files of various legume species, including *Glycine max* Wm82.a4.v1, *Glycine soja* 1.1, *Medicago truncatula* Mt4.0v1, *Phaseolus vulgaris* v2.1, *Lens culinaris* v1, and *Cicer arietinum* v1.0, from the Phytozome database. Additionally, the genome and gene GFF3 files for *Pisum sativum* v1a were sourced from the Plant Bioinformatics facility (https://urgi.versailles.inra.fr/). Collinearity analysis was performed using the MCScanX tool^54^, and the identified syntenic gene pairs were visualized in dual synteny plots using TBtools^55^.

## Acknowledgements

We thank Dr. Justin Renaud (London Research and Development Centre, AAFC) for assistance with the mass spectrometry analysis, Dr. Paul Erhardt from University of Toledo for providing the semi-synthetic glyceollin I standard, and Alex Molnar (London Research and Development Centre, AAFC) for technical assistance. This work was supported by Agriculture and Agri-Food Canada’s Abase grant (J-002364) and the ASC-09 Soybean Cluster Activity #7A (J-002060) to SD.

## Author contributions

S.D. conceived the study; S.D. and P.K. designed the experiments; P.K. and K.K. performed experiments and analyzed the data; T.M. analyzed the chemical data; J.L. prepared WGE; P.K., K.K., N.K. and S.D. wrote the manuscript. All authors discussed the data and made comments on the manuscript.

## Additional information

**Figure S1.**
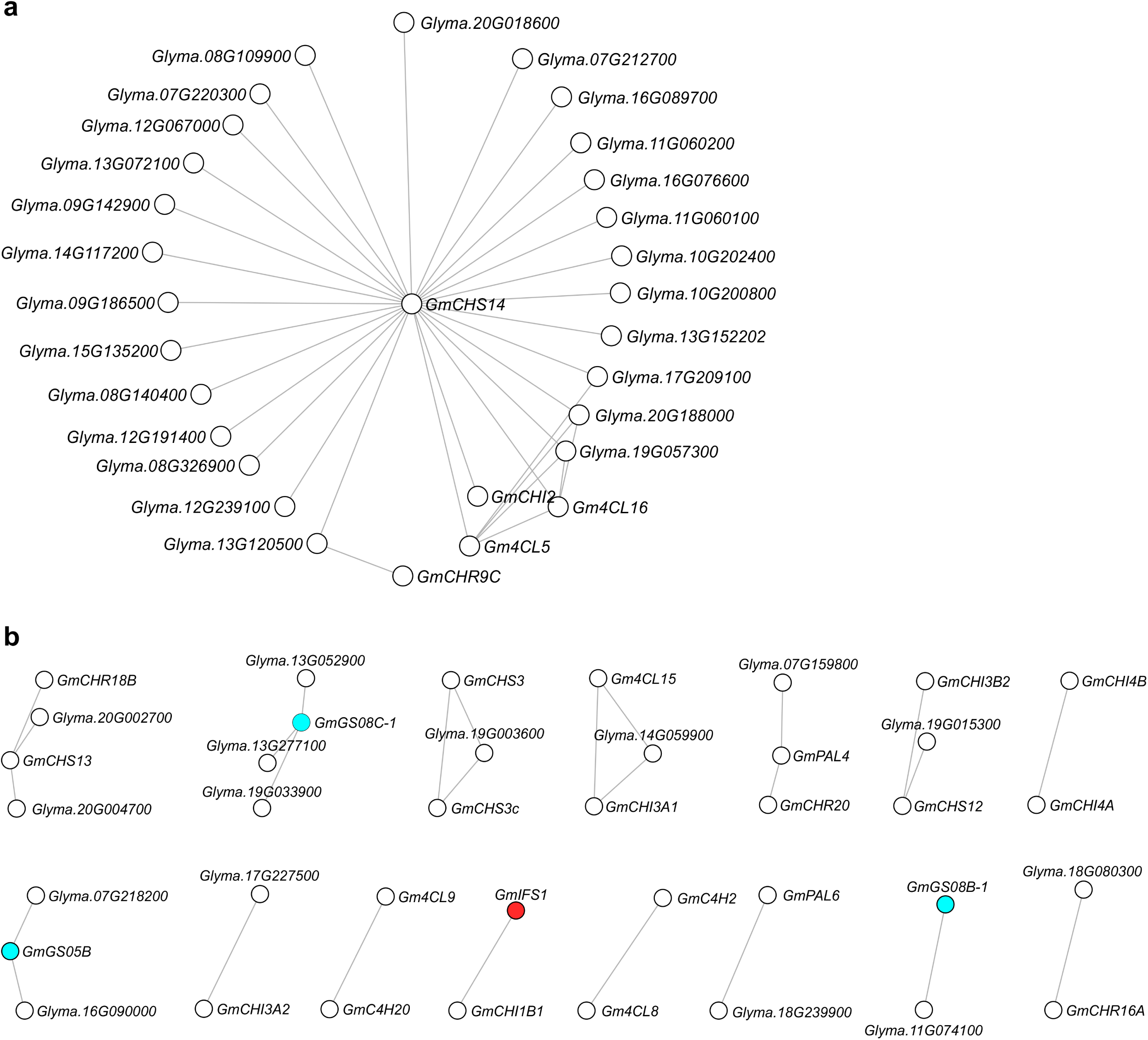
Co-expression networks not included in Fig. 3.

**Figure S2.**
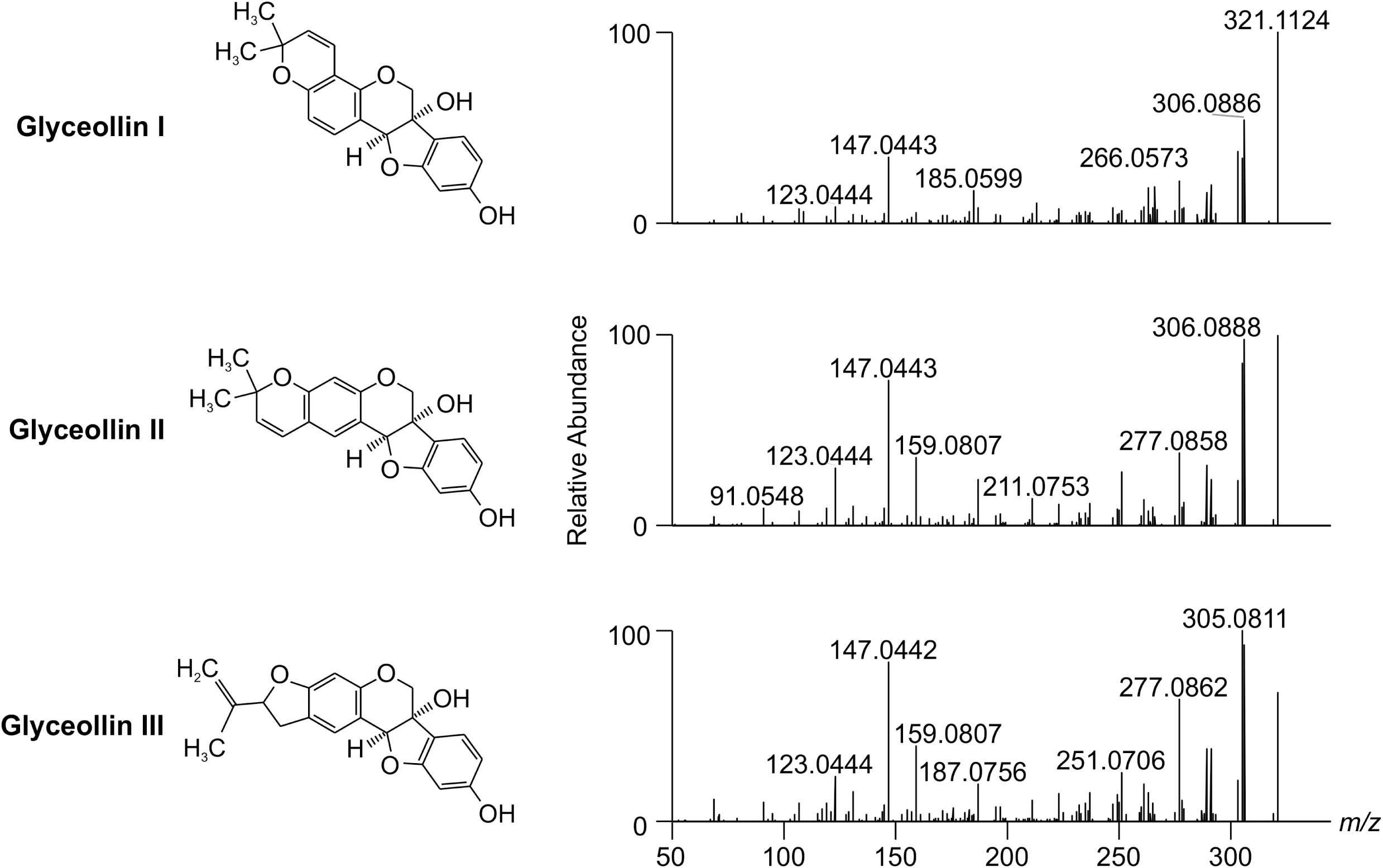
MS2 fragmentation for the three major glyceollins.

**Table S1.** Transcriptome datasets used in the study for identification of *GmGS* gene candidates.

**Table S2.** Collinear_GmGS

**Table S3.** Isoflavonoid genes and locations, transcriptome studies used for co-expression analysis and significant correlation coefficients.

**Table S4.** List of primers used in the study.

