## Supplementary material for "Discovery of the missing cytochrome P450 monooxygenase cyclases that conclude glyceollin biosynthesis in soybean": Table S1

| **Bioproject ID** | **Cultivars used** | **Type of tissue used** | **Sample collection hours post inoculation (hpi)** | **Reference** |
| --- | --- | --- | --- | --- |
| PRJNA324419 | Williams | Hypocotyl | 24 | (Li et al*.*, 2016) |
| PRJNA544432 | Harosoy63 and William82 | Imbibed seeds of Harosoy63 and Hairy roots of William82 | 24 and 48 | (Jahan et al*.*, 2020) |
| PRJNA210431 | Williams and its NILs having Rps1-a, Rps1-b, 1-c and 1-k Rps3-a, 3-b, 3-c, 4, 5, and 6 | Hypocotyl | 24 | (Lin et al*.*, 2014) |
| PRJNA318321 | Williams 82 | Roots | 0.5, 3, 6 and 12 | (Jing et al*.*, 2016) |
| PRJNA478334 | Conrad, Sloan and their RILs | Roots | 24 | (Million et al*.*, 2023) |

**Table S1:** List of high throughput transcriptome bioprojects used in the study.
