## Supplementary material for "Discovery of the missing cytochrome P450 monooxygenase cyclases that conclude glyceollin biosynthesis in soybean": Table S4

**Table S4.** List of primers used for gene expression and gene silencing.

| **Gene** | **Primer Name** | **Sequence (5’→3’)** | **Amplicon size (bp)** |
| --- | --- | --- | --- |
| **Gene Cloning and Expression** | | | |
| *GmGS01A* | GmGS01A-UTR-F | CTACCTAGCTAGCTAGCTAAAATGG | 1,650 |
|  | GmGS01A-UTR-R | ATATAGTTTAACAATCAACAATACACACA |  |
|  | GmGS01A-GW-F | GGGGACAAGTTTGTACAAAAAAGCAGGCTTCATGGAATTAGTTCTACATTTCCTAAAC | 1,581 |
|  | GmGS01A-GW-R-st | GGGGACCACTTTGTACAAGAAAGCTGGGTCTCACATACTTTTGTAACAACTTGGAG |  |
| *GmGS03B* | GmGS03B-UTR-F | GACGAATGGTAAGACTCTAAGGATG | 1,699 |
|  | GmGS03B-UTR-R | ATATACGTGAATGGAAAAATGATGTT |  |
|  | GmGS03B-GW-F | GGGGACAAGTTTGTACAAAAAAGCAGGCTTCATGGCTTTTCAAGTGTTGTTCATTT | 1,533 |
|  | GmGS03B-GW-R-st | GGGGACCACTTTGTACAAGAAAGCTGGGTCTCACACAACAGGGAATGGGTTAAG |  |
| *GmGS05A* | GmGS05A-UTR-F | ACACGTCCATCTCTGGATAACAA | 1,677 |
|  | GmGS05A-UTR-R | TTAACATCAAACAACAGATACAAAGCT |  |
|  | GmGS05A-GW-F | GGGGACAAGTTTGTACAAAAAAGCAGGCTTCATGAAACCCACGGCAACATT | 1,527 |
|  | GmGS05A-GW-R-st | GGGGACCACTTTGTACAAGAAAGCTGGGTCTTATCTACGAACCACTAGGCATCG |  |
| *GmGS05B* | GmGS05B-UTR-F | TCAGCACAATAGTATTAGGTTCAGAG | 1,709 |
|  | GmGS05B-UTR-R | AAAACATAACAAATTAAAGCACCC |  |
|  | GmGS05B-GW-F | GGGGACAAGTTTGTACAAAAAAGCAGGCTTCATGTCTCCTTGGGTTATAGCCG | 1,536 |
|  | GmGS05B-GW-R-st | GGGGACCACTTTGTACAAGAAAGCTGGGTCTCACAATGACAATGATGAATACACAT |  |
| *GmGS08B-1* | GmGS08B-1-UTR-F | CGTACTTAATCTGACCCATAAATAGCA | 1,678 |
|  | GmGS08B-1-UTR-R | AAATCCACCCCAGCAAACATAA |  |
|  | GmGS08B-1-GW-F | GGGGACAAGTTTGTACAAAAAAGCAGGCTTCATGATTTGGATAGCAGCTTTTTTA | 1,497 |
|  | GmGS08B-1-GW-R-st | GGGGACCACTTTGTACAAGAAAGCTGGGTCCTAATCACTTGCAGTGTGAAGCC |  |
| *GmGS08C-1* | GmGS08C-1-UTR-F | GTATACACATACATGCACACTGAAATCT | 1,689 |
|  | GmGS08C-1-UTR-R | TGTGTCCACATTAGCCAAACC |  |
|  | GmGS08C-1-GW-F | GGGGACAAGTTTGTACAAAAAAGCAGGCTTCATGGACAACATTTGGATTGTGT | 1,425 |
|  | GmGS08C-1-GW-R-st | GGGGACCACTTTGTACAAGAAAGCTGGGTCTTAAGATTCCACTCTCCTTACCTTC |  |
| *GmGS08C-2* | GmGS08C-2-UTR-F | TGATCTGAGACCGACATTCCC | 1,530 |
|  | GmGS08C-2-UTR-R | CCACACCCACCTTGTTCTCTT |  |
|  | GmGS08C-2-GW-F | GGGGACAAGTTTGTACAAAAAAGCAGGCTTCATGCTTAAATCAAAACGTAAGAAGAATA | 1,347 |
|  | GmGS08C-2-GW-R-st | GGGGACCACTTTGTACAAGAAAGCTGGGTCTTAAGATTCCACTCTCCTTACCTTC |  |
| *GmGS10A* | GmGS10A-UTR-F | TTAATTTCCCTCCCCGTTTTG | 1,692 |
|  | GmGS10A-UTR-R | GATTTTCTTGCAGCCACACTACAT |  |
|  | GmGS10A-GW-F | GGGGACAAGTTTGTACAAAAAAGCAGGCTTCATGGAGTTTGTAACTTGTGCATT | 1,494 |
|  | GmGS10A-GW-R-st | GGGGACCACTTTGTACAAGAAAGCTGGGTCTTAGTTGCTTATGTTAATAGGAACAAC |  |
| *GmGS11A* | GmGS11A-UTR-F | GACCAGTTGATTCTATTTCCCAT | 1,637 |
|  | GmGS11A-UTR-R | GTAGAGATATCAACAGGTACTACAAACAT |  |
|  | GmGS11A-GW-F | GGGGACAAGTTTGTACAAAAAAGCAGGCTTCATGGAACATTCTCAACTGTCCATTGTCATTA | 1,518 |
|  | GmGS11A-GW-R-stop | GGGGACCACTTTGTACAAGAAAGCTGGGTCTCATGTAGCTTGGTAAACGGTGGGAAT |  |
| *GmGS11B* | GmGS11B-UTR-F | AACTCAAAGGATTCTCAGCAAACA | 1,752 |
|  | GmGS11B-UTR-R | ATTACATCAAACTAGACTCCCCAAAG |  |
|  | GmGS11B-GW-F | GGGGACAAGTTTGTACAAAAAAGCAGGCTTCATGGAATATTCTCCATTGTCCATTGTTATTA | 1,515 |
|  | GmGS11B-GW-R-st | GGGGACCACTTTGTACAAGAAAGCTGGGTCTCATGAAGCTTCATAAACAGTGGGAAT |  |
| *GmGS13A* | GmGS13A-UTR-F | CTCTCTGCTATTTAAACTGATGATGG | 1,674 |
|  | GmGS13A-UTR-R | AAATACAGACAGCAGGTTCACACA |  |
|  | GmGS13A-GW-F | GGGGACAAGTTTGTACAAAAAAGCAGGCTTCATGGAGTTAGTTCTAAACAGCACAACA | 1,569 |
|  | GmGS13A-GW-R-st | GGGGACCACTTTGTACAAGAAAGCTGGGTCTTAGATACTTTCATAACAACTAGGAGATAGGC |  |
| *GmGS13B* | GmGS13B-UTR-F | TGCAACAATTGCAACATAACTA | 1,646 |
|  | GmGS13B-UTR-R | GAACAATAAGGATCACACGACTG |  |
|  | GmGS13B-GW-F | GGGGACAAGTTTGTACAAAAAAGCAGGCTTCATGGACCTTCTCCTAAATTG | 1,584 |
|  | GmGS13B-GW-R-st | GGGG AC CAC TTT GTA CAA GAA AGC TGG GTCTTATAAAGTTTCATAATAGTTG |  |
| *GmGS16A* | GmGS16A-UTR-F | CTATCAACTGAGAAACCCAATGTG | 1,661 |
|  | GmGS16A-UTR-R | CCTTATTTTTCTTTTCAGGTATTTCC |  |
|  | GmGS16A-GW-F | GGGGACAAGTTTGTACAAAAAAGCAGGCTTCATGTGGATCTCTCAGCAAGAAAACTCT | 1,554 |
|  | GmGS16A-GW-R-st | GGGGACCACTTTGTACAAGAAAGCTGGGTCTTATGCATGTGGGGATGCAATT |  |
| *GmGS19B* | GmGS19B-UTR-F | TTAGTTTCTTTCTCTTACCTCACCA | 1,731 |
|  | GmGS19B-UTR-R | ATCACAACATTTTCACCATTCATA |  |
|  | GmGS19B-GW-F | GGGGACAAGTTTGTACAAAAAAGCAGGCTTCATGGCTTATCAAGTGTTGGTGATT | 1,509 |
|  | GmGS19B-GW-R-st | GGGGACCACTTTGTACAAGAAAGCTGGGTCTCACATAACAGGGAATGGGTTAAGCCTTGGA |  |
| *GmGS19C* | GmGS19C-UTR-F | AAAAAGCACCTGATAATTGATAGTA | 1,805 |
|  | GmGS19C-UTR-R | GGAGAGAGCAACAAAAAGCAA |  |
|  | GmGS19C-GW-F | GGGGACAAGTTTGTACAAAAAAGCAGGCTTCATGGCTTATCAAGTGTTGGTGATTTGTG | 1,530 |
|  | GmGS19C-GW-R-st | GGGGACCACTTTGTACAAGAAAGCTGGGTCTCACATAGTAAGGAATGGGTTAATTCTTGG |  |
| *GmGS20A-1* | GmGS20A-1-UTR-F | TAACAGAGATATACATAAAAAAGAGAGAGA | 1,664 |
|  | GmGS20A-1-UTR-R | TTGGGTGAGCTGGACTAAGAG |  |
|  | GmGS20A-1-GW-F | GGGGACAAGTTTGTACAAAAAAGCAGGCTTCATGGAGTTTCTAGGGCTAGCTG | 1,560 |
|  | GmGS20A-1-GW-R-st | GGGGACCACTTTGTACAAGAAAGCTGGGTCCTAAAGGGGCTGAACTATGAGTG |  |
| *GmGS20A-3* | GmGS20A-3-UTR-F | CCGTGAGACAGATCGATACAAAG | 1,225 |
|  | GmGS20A-3-UTR-R | ATATCATTTATATGTCAACAACATACCTTTT |  |
|  | GmGS20A-3-GW-F | GGGGACAAGTTTGTACAAAAAAGCAGGCTTCATGGAGTTTCTAGGGCTAGCTG | 1,146 |
|  | GmGS20A-3-GW-R-st | GGGGACCACTTTGTACAAGAAAGCTGGGTCTCAACAACATACCTTTTTCAAACC |  |
| **Gene Silencing** | | | |
| *GmGS11A* | GmGS11A-hpin-F1 | GGGGACAAGTTTGTACAAAAAAGCAGGCTTCAACTCCTAGGACCAGTTGATTCT | 252 |
|  | GmGS11A-hpin-R1 | GGGGACCACTTTGTACAAGAAAGCTGGGTATAGTGCTACCTGATGAAGGTTAC |  |
| *GmGS13A* | GmGS13A-hpin-F1 | GGGGACAAGTTTGTACAAAAAAGCAGGCTTCCTGAAAATGGATGACCTCTC | 253 |
|  | GmGS13A-hpin-R1 | GGGGACCACTTTGTACAAGAAAGCTGGGTATTGGAGGACCCTCTTCCC |  |
| **qPCR** | | | |
| *GmGS11A* | qGmGS11AF1 | GGCAACAACAGTGAAGCACAGC | 73 |
|  | qGmGS11AR1 | CGAGGCTCCCACTTTGTTGGATTC |  |
| *GmGS11B* | qGmGS11BF1 | CCTCGAGGTTCCAATGACGATG | 149 |
|  | qGmGS11BR1 | ATTCAGCTTGTGCCTTCTCCTTC |  |
| *GmGS13A* | qGmGS13AF1 | AGTTCCATCACGGAGCTCTTCC | 76 |
|  | qGmGS13AR1 | TCAACTCCACAGTGGCAAAGCC |  |
